# An automation workflow for high-throughput manufacturing and analysis of scaffold-supported 3D tissue arrays

**DOI:** 10.1101/2022.08.20.504600

**Authors:** Ruonan Cao, Nancy T Li, Jose L Cadavid, Simon Latour, Cassidy M Tan, Alison P McGuigan

## Abstract

The success rate of bringing novel cancer therapies to the clinic remains extremely low due to the lack of relevant pre-clinical culture models that capture the complexity of human tumours. Patient-derived organoids have emerged as a useful tool to model patient and tumour heterogeneity to begin addressing this need. Scaling these complex culture models while enabling stratified analysis of different cellular sub-populations remains a challenge, however. One strategy to enable higher throughput organoid cultures that also enables easy image-based analysis is the Scaffold-supported Platform for Organoid-based Tissues (SPOT) platform. SPOT allows the generation of flat, thin and dimensionally-defined microtissues in both 96- and 384-well plate footprints and is compatible with tumour organoid culture and downstream image-based readouts. SPOT manufacturing is currently a manual process however, limiting the use of SPOT to perform larger-scale screening. In this study, we integrate and optimize an automation approach to generate tumour-mimetic 3D engineered microtissues in SPOT using a liquid handler, and show comparable within-sample and between-sample variation as the standard manual manufacturing process. Furthermore, we develop a liquid handler-supported whole-cell extraction protocol and as a proof-of-value demonstration, we generate 3D complex tissues containing different proportions of tumour and stromal cells and perform single-cell-based end-point analysis to demonstrate the impact of co-culture on the tumour cell population specifically. We also demonstrate we can incorporate primary patient-derived organoids into the pipeline to capture patient-level tumour heterogeneity. We envision that this automated workflow integrated with 96/384-SPOT and multiple cell types and patient-derived organoid models will provide opportunities for future applications in high-throughput screening for novel personalized therapeutic targets. This pipeline also allows the user to assess dynamic cell responses using high-content longitudinal imaging or downstream single-cell-based analyses.

## 1. Introduction

Drug discovery, particularly in oncology, is challenging because classic pre-clinical models often do not accurately predict the effectiveness of therapies in patients [1]. This is in a large part because conventional 2D cell culture models and simplified spheroid 3D models constructed from immortalized cell lines, typically fail to capture complex and dynamic cellular responses, as these models lack the cellular heterogeneity and microenvironmental cues [2,3] necessary to replicate physiologically relevant cell behaviour. To address this challenge, 3D *in vitro* models that incorporate patient-derived cells or patient-derived organoids (PDOs) have emerged as a powerful strategy to improve pre-clinical assessment accuracy [4–9]. These next-generation engineered models better recapitulate the tumour heterogeneity and enable modelling patient specific disease, but present challenges to culture at scale for high-throughput and automated screening applications [10–12].

The growth of tumour organoids relies on the presence of 3D bio-matrix, such as Matrigel™, which has poor structural integrity that can result in uncontrolled culture deformation when using high cell densities, during long-term culture, or even during the frequent washing steps necessary for analysis [13]. A variety of strategies have emerged to better preserve the fragile 3D cultures including models incorporating a solid substrate such as elastomer-based microfluidic devices [14–16] and scaffold-supported platforms [13,17–22]. By improving the structural integrity and control over culture assembly, some of these models have also enabled increased culture complexity and relevance such as the incorporation of multiple cell types including cancer-associated fibroblasts [13,23], mesenchymal stromal cells [24,25] and endothelial cells [26]. However, most of these novel 3D models are generally low-throughput, require manual manufacturing, and do not integrate easily with standard instrumentation in the existing drug discovery pipeline. Some examples have been reported where automation is beginning to be incorporated into workflows using these complex culture models, primarily with a focus on automation of 3D microtissue/tumour fabrication and/or dispensing drugs to perform a drug response assay [4,6,8,15,27–33]. For example, engineered microtissues/tumours have been fabricated automatically using bioprinting techniques to control the deposition of cell-laden bioinks, such as extrusion-based bioprinting [29,30], stereolithography [31], acoustic bioprinting [32], magnetic force field [8], contact-capillary wicking to infiltrate cell-gel into existing scaffolds [22,34], and FRESH bioprinting using sacrificial support materials [33]. However, the remaining manual downstream processes, such as maintenance, screening and high-content analysis, still significantly limit the throughput of these culture platforms and potentially introduce unwanted experimental variation. In a few cases automation of the entire workflow has been achieved using complex customized systems [35] or integration of multiple different robotic set-ups [36], however, the ease of adapting these approaches for wider applications remains unknown [35]. A need exists therefore for strategies to integrate automation into each step of the fabrication, maintenance, fixation, staining, and high-content analysis of these models.

Another challenge associated with next-generation engineered culture models is the ability to perform single-cell-based analyses in a large-scale [4,6,8,14,35–37]. Moss et al. were able to automate printing, cell seeding, media changes, media collection, and long-term incubation by applying several different robotic set-ups. However, their approach still lacked a single-cell resolution analysis [36]. Bulk cellular read-outs, such as cell viability [4,6,8], protein expression [35,36], permeability [35] and bulk fluorescence [14], offer an accumulated metric of the drug-response from a heterogeneous cell population within each whole-well. This makes it particularly challenging to decipher particular cellular responses from specific sub-populations in a complex microenvironment such as a tumour where multiple cell types are typically present. In another example, Renner et al. reported a fully automated workflow using a standard liquid handler rather than custom equipment to screen neural 3D organoid structures [38]. Automation was integrated throughout the generation, maintenance and downstream analysis pipeline and includes the use of an imaging approach with single-cell resolution [38]. However, this workflow did not involve 3D biomatrix, which could limit the use of this technique for application, such as tumour organoids, where a bio-matrix is critical to maintain cell growth [39]. Further, it is known that stiffness of the culture biomatrix affects tumour cell phenotypes, such as chemosensitivity, therefore incorporating a matrix is likely critical for cancer culture platforms [2,40,41]. The use of a 3D matrix, as is typical in many engineered culture models however, significantly increases the complexity of image-based single cell analysis as these models are typically formed into a thick gel plug geometry with a curved meniscus surface profile. Further the thick geometry of the plug distributes cells into multiple image plane often necessitating the use of confocal microscopy to decipher different cell populations. Microtissues/tumours manufacturing approaches that involve micromolding of the hydrogel matrix to ensure a flat and thin gel geometry can significantly improve the capacity to perform imaging and obtain single cell resolution data [13,22] however to date these systems have not been automated with a view towards scaling throughput.

One micromolding platform that enables the formation of geometrically defined, flat thin, microtissue/tumour cultures is the Scaffold-supported Platform for Organoid-based Tissues (SPOT) [22]. SPOT has been developed in both 96- and 384-well formats and SPOT plates are pre-assembled to enable off-the-shelf use, which make it particularly advantageous for integration with standard automation instrumentation. SPOT has also been shown to enable real-time tracking of patient-derived tumour organoids (**Figure 1A**) [22]. Here, we describe the automation of the manufacturing and analysis workflow for the SPOT model that integrates generation, maintenance, high-throughput treatment, automated microscopy, gel-digestion for cell extraction, and high-throughput flow cytometry of complex 3D microtissues containing patient-derived cells (**Figure 1B**). Our integrated automated workflow removes labour-intensive and challenging microtissue-handling steps using an open-source and inexpensive Opentrons™ OT-2 (OT2) liquid handler, thereby allowing for rapid platform scale-up and implementation into existing screening facilities. Our automated manufacturing and analysis workflow enables *in vitro* culture of highly heterogeneous PDO cell populations combined with bulk phenotypic or single cell analysis at scale.

**Figure 1.**
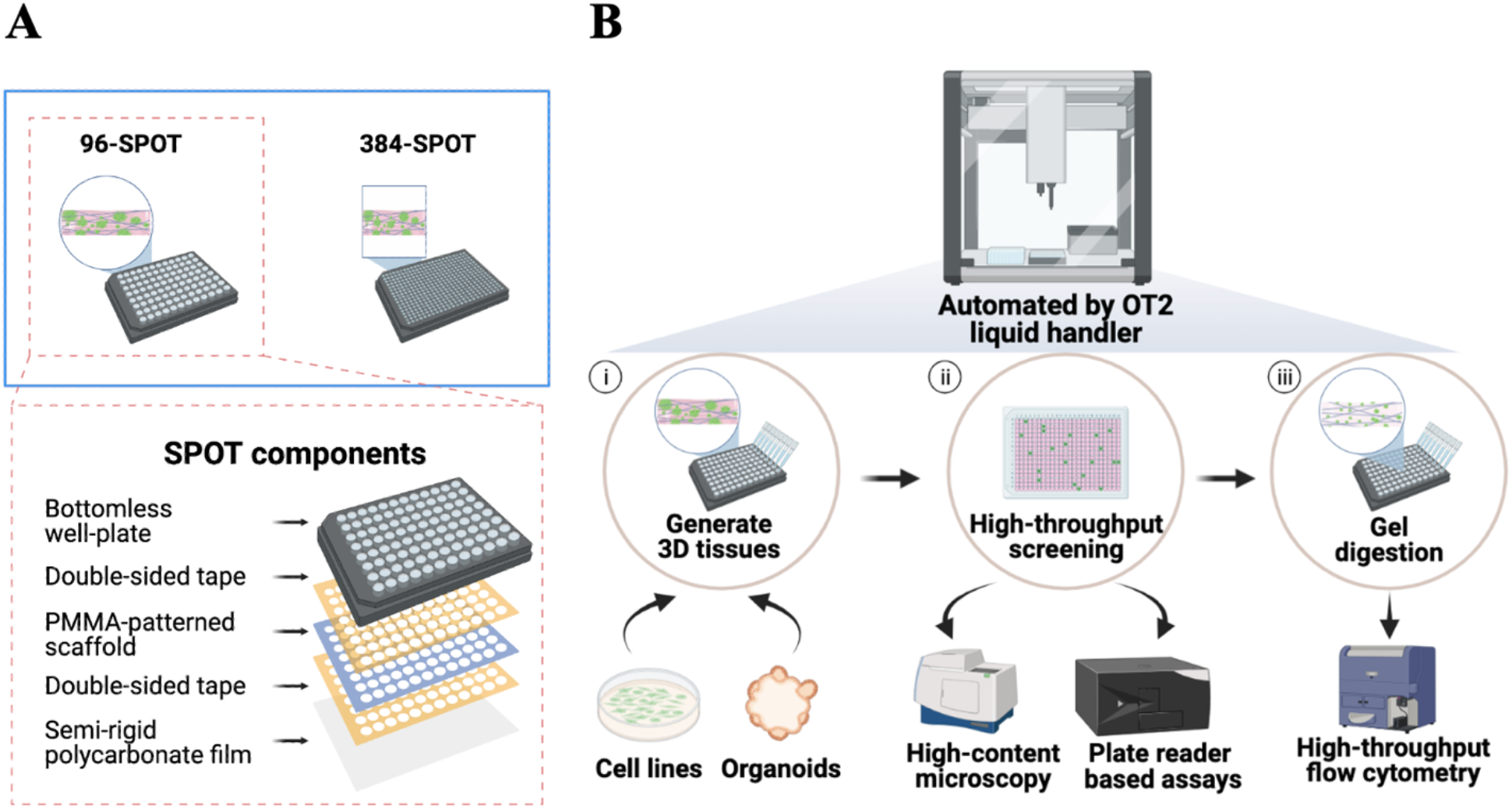
An integrated high-throughput automation workflow to generate, perturb, and analyze 3D in vitro SPOT cultures. (A) Schematic representation of the 96-SPOT components. 96/384-SPOT uses an embedded PMMA-patterned scaffold sheet to support the growth of 3D cell-gel microtissues. The PMMA-patterned paper scaffold (shown in blue) is attached to a bottomless well-plate (top, shown in black) using two layers of double-sided tape (shown in yellow) and a semi-rigid polycarbonate film (bottom). The semi-rigid polycarbonate film is optically transparent and thin to enable microscopy. (B) Schematic representation of the proposed semi-automated workflow performed by the Opentron™ OT-2 (OT2) liquid handler. (i) The OT2 is used to automate the generation of engineered 3D microtissues. Immortalized cell lines or organoid-based cells can be incorporated into SPOT using the optimized cell-gel dispensing OT2 sequence. (ii) The OT2 automates high-throughput screening by assisting in drug and reagent addition and culture maintenance. The engineered tissue arrays can be analyzed by time-course high-content microscopy and plate-reader-based assays, such as AlamarBlue™. (iii) The OT2 is also used to automate gel digestion. The cell-gel within the tissue is mechanically and enzymatically degraded using an optimized digestion sequence. Single cells can then be extracted for end-point downstream analysis, such as high-throughput flow cytometry (bottom).

## 2. Method

### 2.1 Cell culture

The KP4 pancreatic tumour cell line (JCRB Cellbank) was cultured in IMDM containing 10% fetal bovine serum (FBS) (Fisher Scientific, USA), and 1 μg/ml penicillin and streptomycin (Sigma-Aldrich, Canada). The pancreatic stellate cells (PSCs) (ref#3830, ScienceCell) were cultured in DMEM containing 10% fetal bovine serum and 1 μg/ml penicillin and streptomycin. PSCs were used for a maximum of 12 passages. Cells were passaged every three to four days. Pancreatic ductal adenocarcinoma organoids established from PDAC patients were obtained from the UHN Biobank at Princess Margaret Cancer Centre (PMLB identifier PPTO.46 from the University Health Network, Canada) under a protocol in compliance with the University of Toronto Research Ethics Board guidelines (protocol #36107). Organoid cultures were maintained in Advanced DMEM/F-12 (Gibco, ThermoFisher Scientific, USA) supplemented with 2 mM GlutaMAX, 0.5 μM A83-01 (Tocris Biosciences, Bristol, United Kingdom), 10 nM Gastrin I (1-14) (Sigma Aldrich, USA), 10 mM HEPES (Gibco), 1% penicillin/streptomycin, 1X W21 supplement (Wisent), 1.25 mM N-Acetyl-L-cysteine, 10 μM Y-27632 (Selleck Chemicals, USA), 10 mM nicotinamide (Sigma Aldrich), 20% v/v Wnt-3a conditioned media, 30% v/v human R-spondin1 conditioned media (Princess Margaret Living Biobank, Canada), 50 ng/mL recombinant human EGF, 100 ng/mL recombinant human noggin and 100 ng/mL recombinant human FGF-10 (Peprotech, USA). PPTO.46 PDAC organoid cells were cultured in 48-well polystyrene plates in 40 μL domes of Growth Factor Reduced Phenol Red-Free Matrigel™ Matrix (Corning Life Sciences, Corning, USA) with 500 μL of complete media. PPTO.46 PDAC organoid cells were passaged once per week (1:8 split ratio). PDAC organoids were used for a maximum of 40 passages. All cultures were maintained in a humidified atmosphere at 37 °C and 5% CO2.

### 2.2 Lentivirus production and cell transduction

KP4 cells and PPTO.46 were transduced to express green fluorescence protein (GFP), while PSCs were transduced to express blue fluorescence protein (BFP). GFP lentivirus was produced using calcium phosphate co-transfection of HEK293T cells with the psPAx2 plasmid (packaging vector, Addgene #12260), pMd2g plasmid (VSVG envelope, Addgene #12259) and the pLenti-CMV-GFP-Puro plasmid (Addgene #17448) as previously described [42]. BFP lentivirus was produced similarly with pBFP2-IRES-Neo (Addgene #108175). Supernatants containing viral particles were harvested 48 h after transfection and concentrated using a 20 mL 100,000 MWCO spin column (Vivaspin ®, USA). Wild-type KP4 cells were transduced with the concentrated virus, and GFP-expressing cells were selected using puromycin selection (1 μg/mL, Sigma-Aldrich, USA). Wild-type PSC cells were transduced similarly and Neomycin (300 μg/mL, ThermalFisher, USA) was used for selection. GFP-expressing PDOs were created using the same virus as KP4 but with a slightly different transduction protocol. PDOs were suspended in infection media and spinoculated for 1 hour at 600 g at 32 °C, before being resuspended and incubated for an additional 6 hours at 37 °C in 5% CO2. PDOs were then washed and seeded into Matrigel™ with fresh media. 48 hours after infection, cells were sorted for the middle 80% of the GFP+ population.

### 2.3 SPOT component fabrication and assembly

The 96/384-SPOT plates were fabricated as previously described [22]. Cellulose scaffolds were partially infiltrated with a poly(methyl methacrylate) (PMMA, 120KDa, Sigma-Aldrich, USA) solution in acetone (Sigma-Aldrich, USA) to block the inter-well regions using the AxiDraw V3 (Evil Mad Scientists, USA). The PMMA-acetone solution (1.75g PMMA to 7.84mL acetone) was mixed with blue nail polish (Essie) for visibility. For the 96-SPOT, the solution was dispensed through a 20-gauge standard Luer Lock tip (McMaster-Carr, USA) in constant contact with the cellulose scaffold controlled by the Inkscape (https://inkscape.org/) through an AxiDraw extension (https://wiki.evilmadscientist.com/Axidraw_Software_Installation). As the printhead moved and acetone quickly evaporated, a sheet of PMMA that blocks the pores in the scaffold was left behind making it impermeable. For the 384-SPOT, the solution was dispensed through a 16-gauge standard Luer Lock tip (McMaster-Carr, USA) controlled directly using an interactive python API (https://axidraw.com/doc/py_api/#introduction). For each plate, two sheets of double-sided, poly-acrylic adhesive tape (Adhesive Research, ARcare, Catalog # 90106NB) were cut with an array of 96 circular holes or an array of 384 square holes corresponding to the no-bottom 96 well plate (Greiner, Catalog #82050-714) or no-bottom 384 well plate (Greiner, Catalog # 82051-262), respectively, using the CO_2_ Laser Cutter (VLS3.60, Universal Laser Systems, USA) with 50% laser power and 35% speed. One side of the adhesive tape covers was engraved with two lines running across the interwell space between columns 4-5 and 6-7 for 96-SPOT and columns 9-10 and 13-14 for 384-SPOT with 7% laser power and 100% speed to assist in assembly alignment. All components were UV sterilized and assembled in a sterile environment. The double-sided tape was applied to the PMMA-patterned scaffold sheet with the support of PDMS, followed by another layer of double-sided tape on the other side of the scaffold. Then, the sandwiched structure was aligned to the no-bottom well plate with the aid of engraved lines as the middle tape cover was removed first to align and then the two side pieces were removed to adhere fully. Lastly, a thin, transparent polycarbonate film (McMaster-Carr, Catalog # 85585K102) was cut to 110 mm × 75 mm and attached to the bottom of the well plate. The assembled well plates were clamped prior to use to prevent delamination of the layers.

### 2.4 Paper scaffolds physical properties characterization

Scaffold I (Miniminit Products, R10, Scarborough, Canada), the original scaffold used in 96/384-SPOT, scaffold II (Teeli Bag Filter Large Square, Riensch & Held GmbH & Co.KG, Germany), scaffold III (Finum Tea Filters XL, Riensch & Held GmbH & Co.KG, Germany) and scaffold IV (TR Coffee FIL WS 17.0 CR NAT, Twin Rivers Paper Company, USA) were assessed based on physical and physiological properties. Scanning Electron Microscopy (SEM) images were obtained using a Hitachi SEM SU3500 (Hitachi High-Technologies Canada Inc., Canada). Samples were mounted onto carbon-tape coated stubs and gold–palladium sputter coated for 55 s using a Bal-Tec SCD050 Sample Sputter Coater (Leica Biosystems, USA) and then imaged at 10 kVD. The autofluorescence of each scaffold was imaged at the center of a 96-SPOT well fabricated with corresponding scaffolds using an ImageXpress Pico (IXP) (Molecular Devices, USA), a high-content imaging system using both laser-based and image-based autofocus settings (16-bit), with 300 ms exposure acquired at 4X. To quantify the pore area coverage, brightfield images with 10 ms exposure were taken with the IXP microscope. The images were thresholded using OTSU methods on ImageJ to obtain particles and their coverage area. Edge particles and any particles smaller than 890 μm^2^ were excluded from the analysis. The wicking ability of paper scaffolds was assessed on assembled 96-SPOT. The amount of time required for 5 μL of water-based red ink (Ecoline) to cover one well was obtained from observation of video recordings.

### 2.5 Cellulose scaffold seeding

Collagen hydrogel was prepared by mixing eight parts 3 mg/mL type I bovine collagen (PureCol, Advanced BioMatrix) or 6 mg/mL type I bovine collagen (Nutragen, Advanced BioMatrix), with 1 part 10x minimal essential medium (MEM, Life Technologies, Grand Island, USA) by volume and neutralizing to pH 7 endpoint with 0.8M NaHCO3 (Sigma-Aldrich). The solution was kept on ice. Following a standard trypsinization protocol, the adherent cells, KP4 and PSCs, were pelleted by centrifugation (300g, 5 min, 4 °C) and re-suspended in an appropriate volume of 3 mg/ml and 6 mg/ml collagen, respectively, to achieve the desired cell concentration. In the case of organoids, the PPTO.46 cells were collected by mechanical disruption of the Matrigel™ domes followed by a 10 min incubation in TrypLE Express (Gibco) at 37 °C. This process was repeated until mostly single cells were observed under the microscope (a total of 3 times). Cells were pelleted by centrifugation (300 g, 5 min, 4 °C). In accordance with [43] the PPTO.46 cell pellet was re-suspended in an appropriate volume of a hydrogel blend to achieve the desired cell concentration. The hydrogel blend used for PPTO.46 cells comprised 25% Matrigel™ and 75% 3 mg/ml collagen hydrogel, as reported previously [43].

As described previously [22], 70 μL of sterile PBS was added to each inter-well space in the 96-well plate to prevent evaporation while seeding before use, and then the plate was chilled for 30 minutes to ensure no clumping of cell-gel solution during seeding. For manual seeding, a single-channel micropipette (Gilson) was used to deposit 5 μL of cell-gel into the center of each 96-SPOT well by carefully dispensing the gel above the paper surface and gently touching the tip of the pipette to the paper. For the 384-SPOT, a plate membrane (Sigma-Aldrich, cat no. Z380059) was used to cover each column of wells as they are seeded to minimize evaporation. An 8-channel electronic micropipette (Gilson, model no. P8X10M-BC) was used during manual 384-SPOT seeding. The electronic micropipette was used to dispense 2 μL of cell-gel solution in a column of wells similar to the 96-SPOT with cell-gel solution first dispensed into an 8-well PCR strip. Cell-gel solution was prepared similarly for OT2 seeded plates (Opentrons, United States). The OT2 was calibrated based on manufacturer instruction prior to each run. Two temperature modules (Opentrons, United States) were pre-set and maintained at 4 °C to keep the SPOT plate and the 8-well PCR strip cold during the entire operation. A 110 mm × 75 mm × 4.1 mm aluminum plate was customized and used on top of the Temperature Module (Opentrons, United States) to support the thin polycarbonate plate during seeding along with offering better heat conduction. The protocol was designed based on Opentrons Python API V2 (https://docs.opentrons.com/v2/). The cell-gel solution stored in 8-well PCR strips was mixed before seeding and every six depositions to ensure homogenous cell-gel mixture stock using an 8-channel P300 pipette head (Opentrons, United States) with the default pipette speed of 92.86 μL/s. The 8-channel P20 pipette head (Opentrons, United States) was used for both 96/384-SPOT seeding. The cell-gel solution was aspirated and dispensed at 5 μL/s, and the z-axis speed of the pipette head was set to be 1 mm/s during contacting and moving away from the scaffolds. For both 96/384-SPOT, after seeding the final well, the well plate was allowed to incubate for 1-2 minutes on the cold ice pack or temperature module before incubation for 45 minutes for hydrogel polymerization at 37 °C. Then 200 μL or 60 μL of media was added to each 96-well or 384-well SPOT, respectively, after gelation.

### 2.6 Analysis of cell seeding

GFP-expressing cell (KP4 and PPTO.46) fluorescence was used to assess the variation in seeding between and within wells. Widefield images of GFP-fluorescence in each well were acquired at 4X with the IXP. Four or one sites were acquired per 96- or 384-SPOT well, respectively, to capture an entire well bottom, which was stitched automatically by the IXP software. A custom ImageJ script (adapted from [22,43]) was written to measure the mean gray value (MGV) of the GFP signal of 100 randomly selected squares to obtain the standard deviation within one well. Standard deviations were used to calculate the coefficient of variation associated with each well using the following:

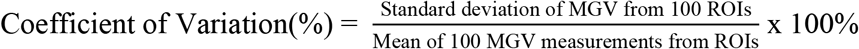

### 2.7 Live dead assay

Wild-type KP4 cells were seeded in 96-SPOT at 5.0 × 10^6^ cells/mL for live-dead assays. 100 μL of 4 μM calcein-AM (Biolegend, the United States) and 4 μg/mL propidium iodide (ThermoFisher, the United States) in IMDM was added to each well and then incubated for 15 minutes at 37 °C. Then, the media was removed, and the well was washed with PBS before adding fresh media. Widefield images of GFP-fluorescence and Texas red-fluorescence in each well were acquired at 4X with the IXP. The MGV was quantified. Similarly, GFP-expressing PPTO.46 cells were seeded in 96-SPOT at 3.0 × 10^6^ cells/mL with 25% Matrigel™ and 75% collagen hydrogel for live-dead assay. 100 μL of 4 μg/mL propidium iodide (ThermoFisher, the United States) in organoid media was added to each well and then incubated for 15 minutes at 37 °C. The GFP fluorescence protein expressed by the cells was used as a surrogate for live-cell stain. The negative dead cell control was generated through incubating 70% ice-cold ethanol for 10 minutes. The wells were then washed and replaced with media for continuous culture.

### 2.8 Quantification of PPTO.46 cell growth in 96-SPOT

The metabolic quantification AlamarBlue™ assay (Sigma), following the manufacturer’s instructions, was used to assess PPTO.46 growth after being seeded by OT2 and compared with the image-based growth metrics. Briefly, GFP-expressing PPTO.46 cells were seeded in 96-SPOT at 3.0 × 10^6^ cells/mL. On days 0, 2, 4 and 6, cells were incubated with 200 μL media with 10% Alamar blue reagent per well for 3 hours as determined previously [22]. Fluorescence intensity was measured at a 3-hour timepoint in a microplate reader at 560/590 nm. Negative control wells containing no cells and positive control wells containing fully-reduced 10% AlamarBlue™ reagent. Images of the different sets of cultures were taken using the IXP on days 0, 2, 4, and 6 as well to visually verify that cell growth was linear over 6-days of culture. The MGV of GFP signal was measured for each microtissue using Image J, showing that the cell growth was linear over 6-days of culture and consistent with the AlamarBlue™ readout.

### 2.9 Gel digestion

The digestion protocol was adapted from a previous protocol [23,44]. Cell rinse buffer was used to dissolve and dilute collagenase XI (Millipore Sigma, USA, Catalog# C7657), DNase I (Sigma-Aldrich, USA, Catalog# D4527) and protease (Millipore Sigma, USA, Catalog# P8811). 100 mL of cell rinse solution was prepared by combining 10 mL 10X modified Earle’s medium, 5 mL of 50mM MgCl2 solution (BioShop, USA, Catalog# 7791-18-6), 83 mL of water, and 2 mL of 1 M HEPES (Thermo Fisher Scientific, Waltham, USA, Catalog# 15630-080) in a sterile 500-mL bottle. The 10X modified Earle’s medium consists of 68 g of NaCl (BioShop, USA, Catalog# 7647-14-S), 10 g of glucose (Sigma-Aldrich, USA, Catalog# 50-99-7), 4 g of KCl (BioShop, USA, Catalog# POC 308), and 1.22 g of anhydrous NaH2PO4 (Sigma Aldrich, USA, Catalog# RDD007) in 930 mL of water. Then the pH was adjusted to 7.3 by titrating with NaOH solution (Sigma-Aldrich, USA). 12mL of digestion solution was prepared by combining 9 mL of cell rinse solution with 1 mL of 2000 U/mL collagenase XI, 3000U/mL DNase I and 3mg/mL protease. The 96-SPOT wells were washed once with PBS and then incubated with 200 μL of digestion solution per well for 45 minutes at 37 °C. To stop the reaction, 20 μL FBS was added per well. Each well was pipetted vigorously using an 8-channel pipette manually or by OT2 using an 8-channel P300 pipette head at 275 μL/s to dissociate cells from gel. After transferring cell suspension to a separate well plate, the wells were rewashed with 200 μL 3% BSA in PBS to harvest most of the cells. Cells were spun to remove supernatant at 50 μL/s for subsequent fixing in 4% PFA for 15 minutes. After two runs of PBS washes, the cells were counted or stored at 4 °C for further analysis.

### 2.10 Measurement of cell proliferation using the EdU assay

A total concentration of 1.5 × 10^7^ cells/mL was seeded into the 96-SPOT with various cell compositions: GFP-expressing KP4 monoculture control, BFP-expressing PSCs monoculture control, and GFP-expressing KP4 and BFP-expressing PSCs cocultured with ratios of 7:3, 1:1, 1:9, and 1:19. All the conditions were cultured in DMEM for three days before performing the EdU assay. 100 μL DMEM media from each well was discarded and replaced with 100 μL DMEM with 20 μM EdU (Abcam, UK, Catalog# ab219801) to reach a final concentration of 10 μM EdU. After 3 hours of incubation at 37 °C, cells were digested out and fixed using the OT2 as described above. Adapted from the manufacturer’s instruction (Abcam, UK), the supernatant was removed from fixed cells using the OT2 at 50 μL/s after spinning at 800 x g at 4 °C for 5 minutes. 100 μL 1X permeabilization buffer (Abcam, UK, Catalog# ab219801)) was added to each well and incubated for 15 minutes at room temperature and then 100 μL of 3% BSA (Sigma-Aldrich, USA) was added into the well followed by spinning and removing the supernatant. 125 μL EdU reaction solution was added per well, which consisted of 438 μL PBS, 10 μL 100 mM CuSO_4_ (Abcam, UK, Catalog# ab219801)), 2.5 μL of 500 μM Texas Red azide (AAT Bioquest, USA) and 50 μL 20 mg/mL sodium ascorbate (Abcam, UK, Catalog# ab219801)) per 500 μL solution, and incubated for 30 minutes at room temperature. Then the cells were washed twice with 3% BSA with the assistance from the OT2 using similar setting as before. Next, cells were re-suspended and analyzed directly from a V-bottom well plate (ThermoFisher, USA, Catalog# 249570) using a LSR Fortessa flow cytometer (BD) coupled to a high-throughput sampler. KP4 and PSC cells were gated based on GFP and BFP fluorescence, respectively, and EdU+ cells were determined as a percentage for each cell population. Cytometry data were analyzed using FlowJo (version 10.8.1).

### 2.11 Statistics

Statistical analysis was carried out in GraphPad Prism 9 (GraphPad Software, USA). Ordinary one-way ANOVA was used for assessing the physical properties of paper scaffolds, basic parameters of OT2, and reproducibility of microtissues seeded by the OT2 sequence. A Student t-test was used for assessing manual and OT2-seeded KP4 microtissues or PPTO.46 microtissues. p < 0.05 was considered significant.

## 3. Results and discussion

### 3.1 Comparison of scaffold candidates to identify material compatible with automation

The overall goal of this study was to establish an automated pipeline that can dispense a cell-gel mixture into 96/384-SPOT to generate 3D tissue arrays, assist in medium/high-throughput screening assays, and digest hydrogel to recover single cells from the SPOT platform for end-point analysis (**Figure 1A**). The 96/384-SPOT utilizes a PMMA-patterned cellulose scaffold and capillary wicking to form thin microgel tissue without a meniscus. Further the cellulose scaffold provides structural support to the hydrogel to improve handlability. The scaffold is attached to a commercial bottomless well plate and to a thin (0.127 mm) polycarbonate film, which acts as a plate bottom, using 2 layers of double-sided tape to allow high-quality imaging and compatibility with plate-reader-based assays overtime (**Figure 1B)**. To provide automated manufacturing and microtissue processing, we utilized the liquid handling automated pipetting system Opentrons™ OT-2 Lab Robot (OT2). This system is open-source, commercially available, and allows for additional attachments for workflow customization, i.e. the HEPA filter module and temperature-controlled modules allow the possibility for a sterile work surface and for dispensing of temperature-sensitive hydrogels, respectively. Initial pilot studies identified that manual dispensing provided flexible control of the pipette tip contact angle with the scaffold during the contact wicking manufacturing process, while the OT2 automated pipetting system allowed only perpendicular dispensing of the cell-gel mix. Perpendicular dispensing led to unsatisfactory gel distribution into the paper scaffold and uneven seeding during manufacturing. To address this design restriction, we therefore first had to select a paper scaffold with faster and more-reproducible capillary wicking properties to ensure consistent cell-gel mixture spreading within the well during automated manufacturing.

To this end, we characterized three scaffold candidates (II, III and IV) and compared them with the original paper scaffold (I) used in 96/384-SPOT [22]. The three scaffold candidates were preliminarily selected from a larger pool of candidates based on thickness (similar thickness as scaffold I), material types (cellulose scaffolds are preferred), and material size compatibility with well-plate dimensions. Scanning electron microscopy (SEM) images of each scaffold (**Figure 2A**) revealed a similar porous structure across all three scaffolds except scaffold IV, which exhibited a more compact cellulous fiber structure. To quantify the capillarity wicking ability, we measured the time it took for color dyed PBS to cover one 96-SPOT well fabricated from one of the four different scaffolds. As shown in **Figure 2B**, scaffold III offered the most superior and consistent wicking ability across multiple wells and sheets of papers. We speculate that these observations may arise from differences in hydrophobicity of each scaffold candidate. Using widefield microscopy images we also calculated the pore area coverage, which further confirmed that scaffold IV had a significantly smaller and lower number of pores: specifically scaffold IV had ~10% lower pore area coverage compared with scaffold I (**Figure 2C**). Scaffolds II and III had a similar physical structure, but exhibited a significant lower pore area coverage of 4% compared with the original paper scaffold I. To assess how the pore coverage difference affect seeding and hence tissue homogeneity, we seeded GFP-expressing KP4 cells suspended in 3mg/mL bovine collagen manually and assessed cell distribution just after seeding or three days later. All four scaffolds allowed KP4 cells growth without spatial restraints (**Figure 2D**). However the spreading of cells in the scaffold IV appeared not as homogenous as the other three scaffolds (I,II and III) (**SI Figure 1A**). To quantify the tissue homogeneity obtained with the different scaffolds, we measured the coefficient of variation, which is a surrogate for tissue homogeneity, using a previously described method [22]. The coefficient of variation provides a measure of the intrawell variation by quantifying the ratio of the standard deviation of 100 randomly selected squares’ mean gray value within the well over the whole-well mean gray value. Our measurements confirmed that the use of scaffold IV led to poor homogeneity, whereas better homogeneity was observed with both scaffold II and III compared to scaffold I (**SI Figure 1B)**. These results indicated that scaffold III was likely the most compatible candidate with OT2 automated cell-gel deposition based on the wicking properties.

**Figure 2.**
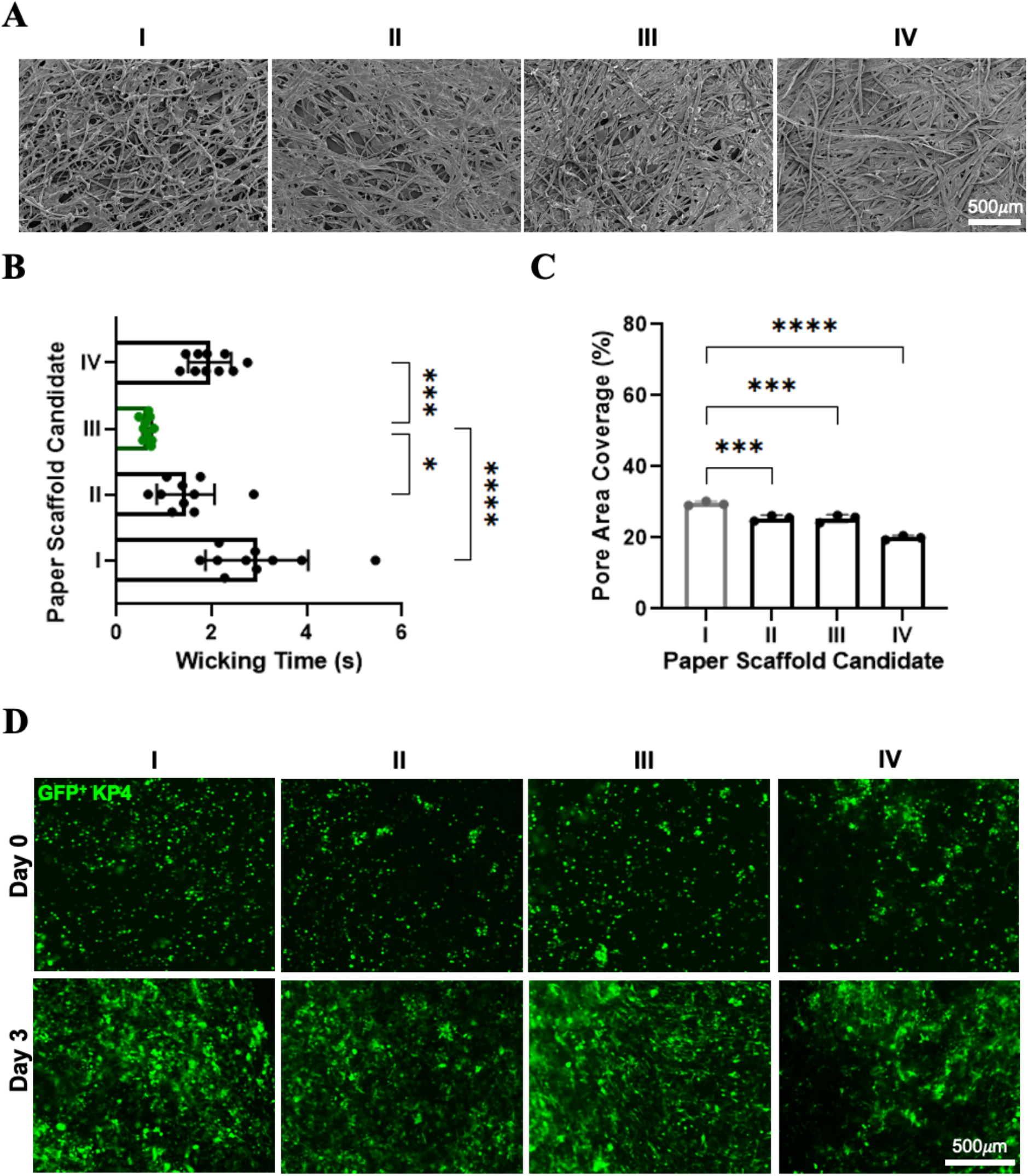
Comparison of scaffold candidates to identify material compatible with automation. (A) Representative SEM images of the original SPOT scaffold (I) and three paper scaffold candidates (II-IV). The scale bar is 500 μm. (B) Wicking time was measured by observing the amount of time required for dyed PBS to wick radially within one well in 96-SPOT for each scaffold candidate. Scaffold candidate III offers the most superior wicking ability (shown in green). Statistical significance was assessed using ANOVA. Mean ± SD of 3 independent experiments. (C) Pore area coverage as a percentage for all four paper scaffolds was measured based on brightfield images taken under 4x magnification. Candidate IV exhibited the most condensed cellulose fibres. Mean ± SD of 3 independent samples size of around 12 mm^2^. (D) Representative images showing GFP-expressing KP4 cells on Day0 and Day 3 seeded at 5 × 10^6^ cells /mL hydrogel using 3mg/mL type I bovine collagen into each scaffold candidate. The scale bar is 500 μm.

One significant advantage of 96/384-SPOT is the ability to easily track cell growth and assess drug treatment effects over time using a widefield microscope. We therefore also assessed whether the new scaffold were compatible with imaging and did not exhibit strong autofluorescence in classically used imaging channels (**SI Figure 2A**). Scaffolds II and III offered lower autofluorescence across DAPI, FITC, Cy3 and Texas Red channels tested compared with scaffold I when they were all exposed for 300 ms (**SI Figure 2B**). We note that scaffold IV could potentially impede high-quality real-time imaging due to its strong autofluorescence. As scaffold III offered a similar porous structure, faster capillary wicking, lower autofluorescence, and supported mammalian cell growth, we selected this scaffold for automated 96/384-SPOT for further optimization.

### 3.2 Characterization and optimization of essential parameters for automated cell-gel dispensing

Having selected a scaffold with appropriate wicking properties, we next set out to identify appropriate liquid handling parameters to enable generation of homogenously seeded scaffolds. Since type I bovine collagen gel, the ECM that has been used previously in the SPOT system, is temperature sensitive, a temperature control setup is needed to prevent the uneven gelation and clumping of cells during seeding. To achieve this, as previously described [22], cold PBS was added into the spaces between the wells of the 96-SPOT to enable temperature and humidity stabilization across the well plate. Moreover, the cell-gel mixture stock solution and the 96/384-SPOT were kept on a temperature-controlled module at 4°C during the gel dispensing process in the OT2 system. To ensure direct contact between the surface of the temperature-controlled module and the bottom surface of the well plate, a customized aluminum block was fitted underneath the plate, providing both improved temperature control and additional physical support for the polycarbonate film during seeding.

As the cell-gel mixture is much more viscous than aqueous solutions (for which the OT2 system was designed for), several parameters needed to be optimized to ensure reproducible small volume deposition into the scaffold. First, we assessed different contact locations of the 8-channel pipette tips and scaffold along the z-axis using a 96-SPOT set up. The plate bottom (z = 0 mm) location was determined based on a regular well-plate thus optimization was required for the SPOT system as it is an off-the-shelf platform. For different tip z-locations we assessed cell-gel infiltration of GFP-expressing KP4 cells using fluorescent images. Specifically, the OT2 tips were lowered in increments of 0.5mm from the default plate-bottom position and dispensed 5uL cell-gel mixture into each well. Due to the viscosity of the cell-gel mixture and interfacial tension on the tips, insufficient or completed failed deposition could occur which led to partially seeded or empty wells. Surprisingly, even small differences in z-increment, introduced a significant difference in the number of empty wells (no GFP KP4 cells observed) for each 8-channel deposition across three z-location settings tested (**Figure 3A(i)**). Dispensing the cell-gel mixture at the plate bottom (z = 0 mm) offered the worst results as most wells contained no cells due to unsuccessful or incomplete contact with paper scaffold. Notably, lowering the tip in the z-direction by 0.5mm versus 1mm showed no significant difference in the number of empty wells observed. However, the tips were noticeably bent with the 1mm setting. Since 96/384-SPOT uses a thin layer of semi-rigid polycarbonate film to facilitate high-quality imaging and plate-reader-based assays, we hypothesized it was essential to prevent denting of the film during the automated seeding process. More importantly, denting the film prohibited us from being capable of auto-focusing which is required for automated imaging. Therefore, we selected a z-displacement for the multichannel tips of 0.5mm below the default plate bottom z-location.

**Figure 3.**
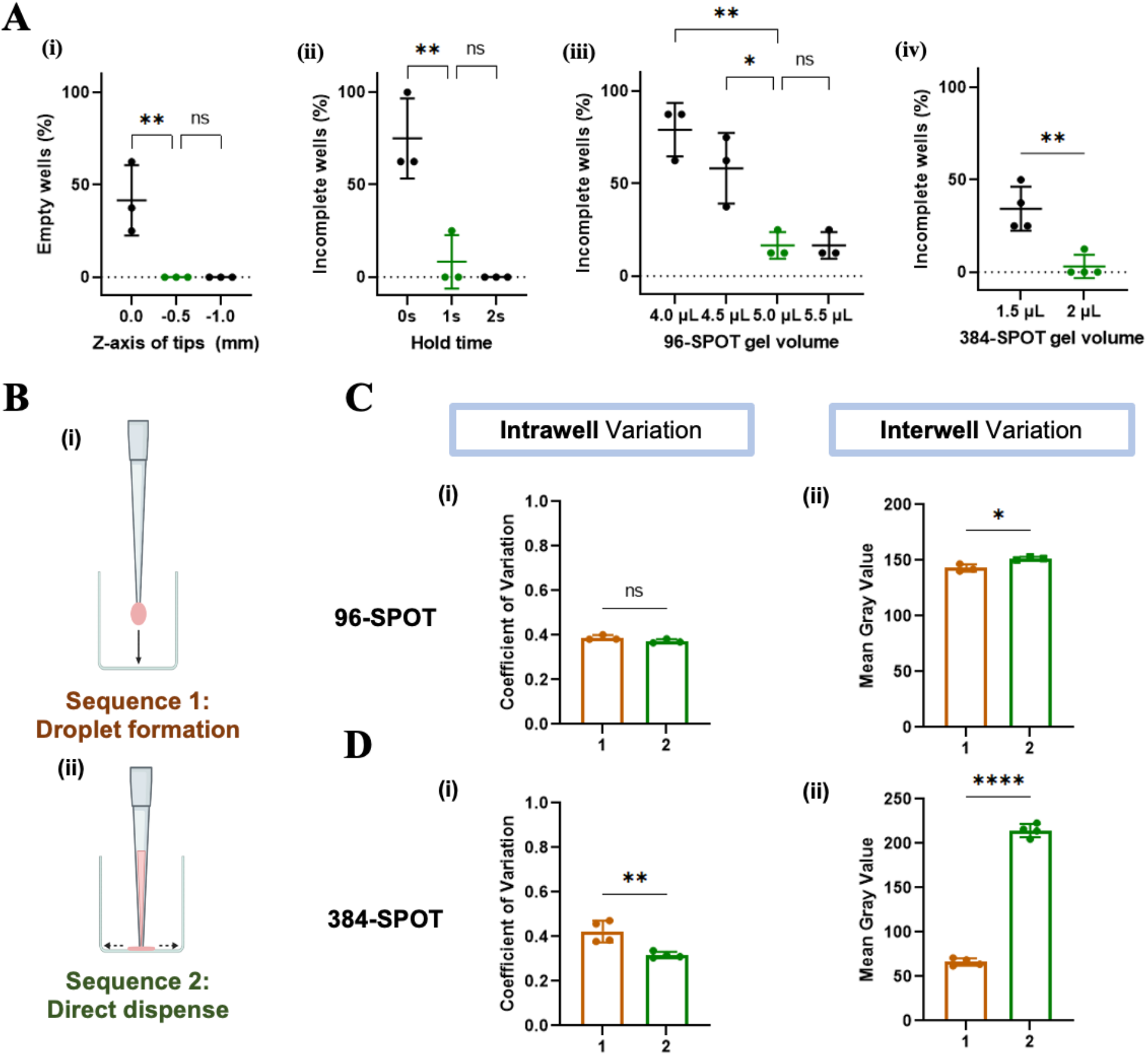
Optimization of OT2 parameters to achieve robust seeding in 96/384-SPOT. (A) GFP-expressing cells were seeded at 30 × 10^6^ cells /mL hydrogel using 3mg/mL type I bovine collagen to evaluate the coverage of cell-gel mixture inside the wells seeded using various parameters. The offset of tip z-axis (i), tip hold time on the paper scaffold after dispensing (ii), 96-SPOT gel volume (iii) and 384-SPOT gel volume (iv) were selected as −0.5mm, 1s, 5mL and 2mL, respectively, for further OT2 sequence optimization. Statistical analysis was performed using ANOVA. (B) Schematic representation of two OT2 cell-gel deposition sequences (i) sequence 1: droplet formation, in which the cell-gel droplet (pink) is formed before contact with the scaffold, and (ii) sequence 2: direct dispense, in which contact is made before the cell-gel (pink) is dispensed. (C) Assessment of (i) intrawell variation using coefficient of variation and (ii) intrawell variation using mean gray value for 96-SPOT. (D) Assessment of (i) intrawell variation using coefficient of variation and (ii) intrawell variation using mean gray value for 384-SPOT. The #2 direct dispense sequence offers consistently better seeding results, especially for 384-SPOT. Statistical analysis was performed using the t-test. Mean ± SD of 3 independent experiments.

Next, as it takes more time for a viscous liquid to travel within the pipette tips due to higher interfacial tension, another parameter that needed to be optimized was the holding time of the pipette tip after the initiation of liquid dispensing. We observed that an additional holding time of 1s after dispensing improved the cell-gel spreading within the well significantly by decreasing the number of incompletely covered wells (**Figure 3A(ii)**). Since further increasing the holding time did not significantly improve dispensing results, we decided to apply 1 s holding time for both 96- and 384-SPOT to prevent clumping of gels inside the tips. We note however that this holding time was optimized for 3mg/mL type I bovine collagen gel, and further optimization might be needed for bio-matrices with much higher viscosity.

Finally, since we selected a different scaffold and different dispensing equipment compared to the original SPOT device, we assessed the minimum volume of gel required to achieve complete infiltration of the well in the SPOT plate. Previously, with manual seeding, 5 μL and 1.5 μL cell-gel mixture were used for 96/384-SPOT, respectively. Using the same cell-gel materials, we tested a range of seeding volumes and imaged the wells. By quantifying the amount of incompletely seeded wells present using each volume, we determined that with the new scaffold material and OT manufacturing process, using a seeding volume of 5 μL and 2 μL for 96/384-SPOT offered optimal and consistent coverage while still preserving cell materials, which is crucial for precious primary samples (**Figure 3A(iii)&(iv)**).

With the basic OT2 parameters optimized, we next set out to identify an appropriate dispensing sequence compatible with 96/384-SPOT. Since the cell-gel mixture is kept at 4°C during the seeding process, we aimed to achieve consistent and reproducible seeding results while minimizing the run time to maintain high cell viability. We extensively assessed two dispensing sequences: #1 droplet formation and #2 direct dispense (**Figure 3B**). Note that we also tested in pilot studies other dispensing sequences, such as using the *blow_out()* function, which pushes an extra amount of air after dispensing liquid, however, none of them were optimal as they either produced extra bubbles or extended run time without a noticeable improvement in seeding homogeneity. The first sequence begins with droplet formation herein called “sequence #1”, which most closely mimicked our manual seeding process, where the cell-gel is first dispensed from a single-channel pipette above the scaffold in order to form a droplet and then the tip is lowered to contact the paper scaffold to initiate wicking. In contrast, to directly dispense, herein called “sequence # 2”, the tip fully contacts the paper scaffold prior to dispensing the cell-gel, and then the tip is slowly lifted away from the scaffold to allow wicking. We compared the intrawell variation (within well) associated with each sequence using the coefficient of variation as described previously while whole-well mean gray value was used to assess the interwell variation (well to well). We found that sequences #1 and #2 offered a similar coefficient of variation for 96-SPOT, indicative of a comparable uniform spreading of the cell-gel mixture within each well (**Figure 3C(i)**). Sequence #1, however, resulted in slightly worse gel deposition into the scaffold as we observed a slightly lower but consistent mean gray value across wells (**Figure 3C(ii)**). Interestingly, we observed a more drastic intrawell difference between the dispense sequences in 384-SPOT, likely due to the small volume used (i.e. 2 μL). Further, sequence #1 failed to deliver an adequate amount of gel into the scaffold, demonstrated by a low mean gray value consistently (**Figure 3C(iii)&(iv)**). We speculate that this was due to the use of such small volumes in which the formed droplet tended to distribute along the pipette tip, potentially due to the interfacial tension. This was not an issue with manual seeding on 384-SPOT since the user could establish an angle between pipette tips and paper scaffold ensuring the entire gel volume wicked into the scaffold. In contrast, when the tip was perpendicular to the scaffold during automated seeding this was not possible. Based on these observations, we selected to move forward with sequence #2 as direct dispensing of the cell-gel offered optimal seeding for both 96/384-SPOT.

### 3.3 Automatically generated microtissues offer similar homogeneity and cell survival as microtissues created using manual seeding

We next set out to compare our optimized OT2 seeding sequence to manual seeding performed by an experienced user. To be robust and reliable, a screening platform needs to exhibit low variation both within a well (intrawell) and between wells (interwell). To measure the robustness of these two features, as described above, we used two image-based metrics, coefficient of variation (intrawell variation) and whole-well mean gray value (interwell variation), to quantify GFP-expressing KP4 cells distributed within each well of 96/384-SPOT on day 0. We observed that seeding of both 96-SPOT and 384-SPOT with OT2 resulted on a lower or at least similar coefficient of variation (intrawell variation) compared to manual seeding (**Figure 4A(i)&(ii)**). Notably, the interwell variation was similar between OT2 seeding and manual seeding (**Figure 4A(iii)&(iv)**). The improvement in intrawell variation associated with OT2 seeding could be explained by more precise and consistent control of the pipette movement when using OT2 compared to manual seeding.

**Figure 4.**
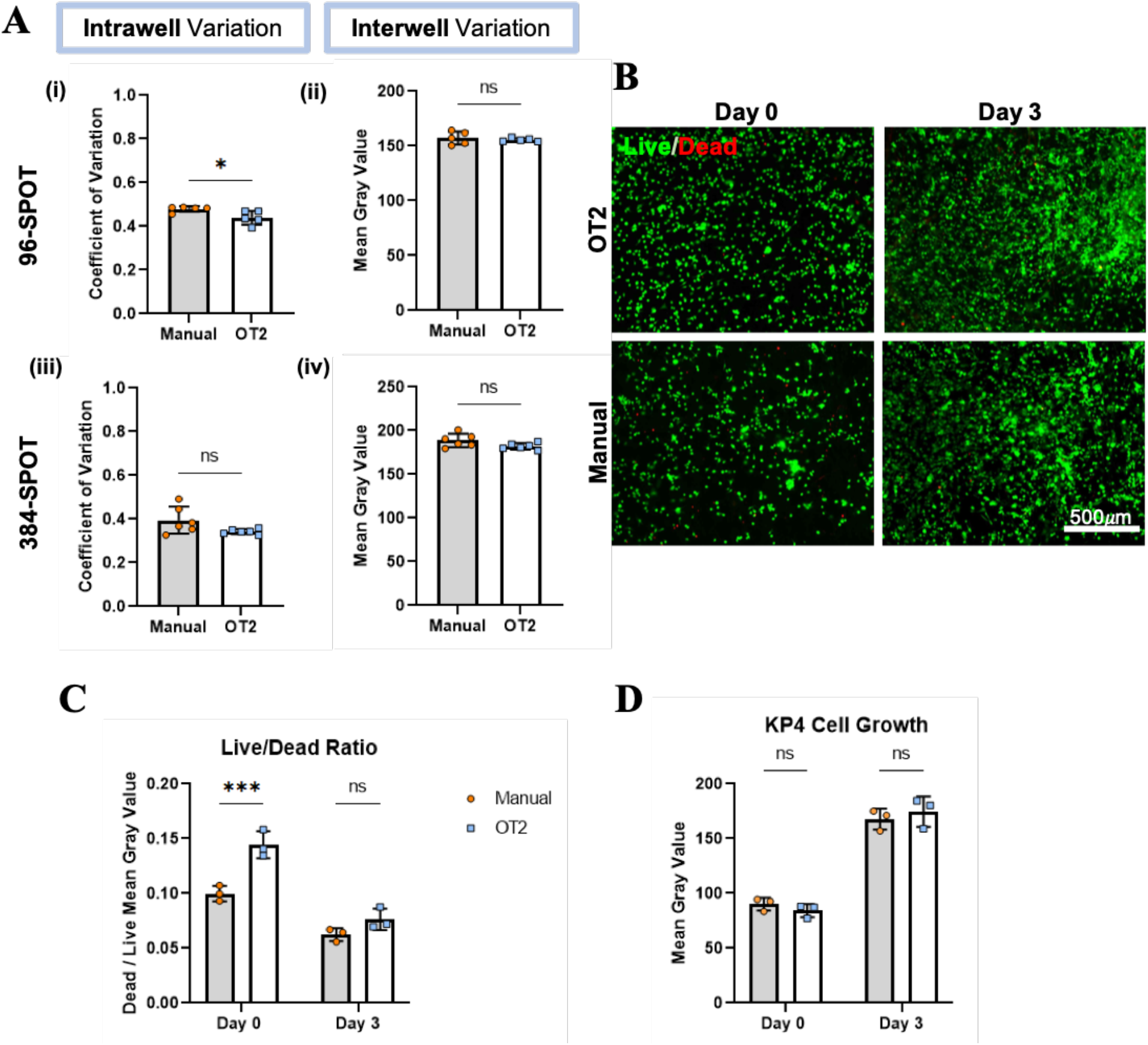
Benchmarking optimized OT2 automated seeding to manual seeding. (A) GFP-expressing cells were seeded at 30 × 10^6^ cells /mL hydrogel using 3mg/mL type I bovine collagen into 96/384-SPOT either by the optimized OT2 sequence or manually by an experienced user. 96-SPOT seeded by the OT2 offered a significantly lower intrawell variation. A similar interwell variation was observed in 96-SPOT using both automation and manual seeding. 384-SPOT seeded by the OT2 had similar intrawell and interwell variations. Statistical analysis was performed using a Student t-test. (B) Representative images of Live/Dead staining using calcein-AM and propidium iodide on Day 0 and Day 3 for OT2 and manual seeded cells. (C) The proportion of dead cells was estimated by obtaining the ratio of dead cells’ mean gray value over live cells’ mean gray value. The OT2 seeded wells showed more cell death only on Day 0. (D) When focusing on the live cell population only, cells seeded by manual and OT2 demonstrated a similar growth pattern characterized by the mean gray value of live cells. Statistical analysis was performed using a Student t-test. Mean ± SD of 3 independent experiments.

While our analysis suggested our OT2 seeding sequence could produce even cell seeding we also wanted to confirm that the automated dispensing sequence did not cause dramatic cell death and hinder cell growth. To do this we assessed cell viability using calcium-AM (staining live cells) and propidium iodide (staining dead cells) cell stains, on both Day 0 and Day 3 for OT2 and manually seeded cells (**Figure 4B)**. We measured the ratio between the mean gray value of dead cells over live cells and compared this metric (i.e. Live/Dead Ratio) between manual and OT2-seeded SPOT microtissues. We found that OT2 seeding did introduced a significantly higher percentage of dead cells at Day 0 (**Figure 4C**). We speculate that this difference may result from the stress experienced by the cells during seeding using the OT2 as the tips are perpendicularly in contact with the paper scaffold which leads to a temporarily high internal pressure inside the tips. However, the difference in cell death between scaffolds seeded manually versus using the OT2 became negligible on Day 3. We also measured cell growth over time to verify that OT2 seeding did not affect the growth kinetic of the seeded cells. To this end, we measured the mean gray value of tissues seeded with OT2 or manually after either 0 days or 3 days and observed no differences at any time point between both the OT2 and manual methods (**Figure 4D**).

### 3.4 Automated dispensing sequences are reproducible in 96/384-SPOT

Having finalized the various parameters for the OT2 we next wanted to assess the reproducibility of our automated OT2 seeding sequence for seeding entire plates. To do this we seeded three entire plates of 96/384-SPOT respectively using the OT2 and quantified heterogeneity using our image-based metrics. Note that after the initial mixing steps, we added an additional mixing step of the cell-gel stock containing the GFP-expressing KP4 cells every six depositions to prevent cell pellet formation at the bottom of the tube. We acquired widefield images after 45 minutes of gelation and the addition of media. The representative images of a entire seeded plate of 96/384-SPOT are shown in **Figure 5A-B**, respectively. By characterizing intrawell (within-well) variance using the coefficient of variation across 3 independently assembled and seeded plates, we found no significant differences among rows or columns using one-way ANOVA for both 96- and 384-SPOT (**Figure 5C-D**). Similarly, we observed no significant difference in interwell (well-to-well) variance based on mean gray value measurements of the whole well for both SPOT plates (**Figure 5E-F**). We were therefore confident that the optimized OT2 sequences could offer reproducible seeding results comparable to those achieved using manual seeding (ref Nancy’s paper).

**Figure 5.**
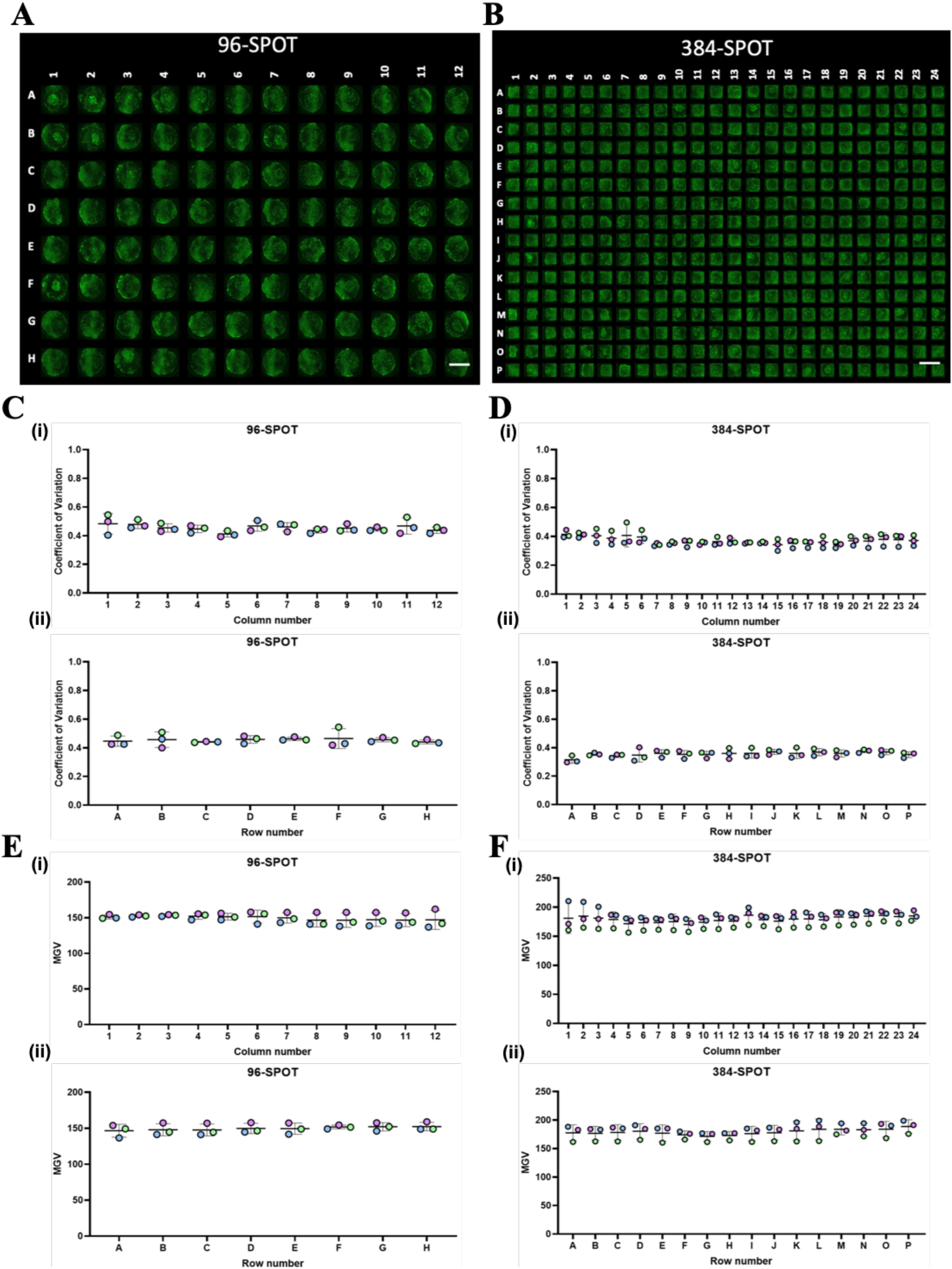
Assessment of fabrication variation associated with 96/384-SPOT generated using optimized OT2 sequence (A) Representative widefield image showing 96-SPOT and (B) 384-SPOT seeded using the optimized OT2 sequence with GFP-expressing KP4 cells (green) at 30 × 10^6^ cells /mL. The scale bar is 5mm. (C) Assessment of coefficient of variation measured from widefield images of 96-SPOT on day 0. Mean ±SD of 3 independent experiments are organized by columns (i) and rows (ii) for 96-SPOT (N=3). ANOVA revealed no statistical significance between columns or rows across 3 independent experiments. (D) Similarly consistent and reproducible results were obtained for 384-SPOT in columns (i) and rows (ii). (E)&(F) The interwell variation for both 96/384-SPOT was quantified based on the mean gray value. No statistically significant differences were observed between columns (i) and rows (ii). Statistical analysis was performed using ANOVA. Mean ± SD of 3 independent experiments.

We also assessed the speed of our workflow to ensure it was within a range to feasibly generate multiple 96 or 384-SPOT plates at once to facilitate larger-scale high-throughput screening. We found it took less than 4 minutes to seed one 96-SPOT and no more than 12 minutes to seed one 384-SPOT, including necessary gel mixing steps between dispensing to ensure a homogenous cell-gel stock. These timeframes are typically our targets when we use a manual seeding process however in reality this seeding pace presents a major logistical challenge for the manual process when attempting to seed a full plate and not simply a few rows within a plate. Further, the OT2 automated seeding enabled consistent run-time from batch to batch, which we anticipate produced more consistent cell viability across plates. Also, the OT2 automated seeding allowed for more consistent well to well seeding run-time which is impossible to achieve through manual seeding. Previously, 384-SPOT manual dispensing was performed using an electronic 8-channel pipette to minimize seeding time and variability. However, achieving optimal seeding consistently using this approach was very challenging for even an experienced user, in part because of the small well size and the risk of misplacing cell-gel droplets to the side of the well. Since the gel dispensing location can be finely controlled by the OT2, the ease of use of the 384-SPOT at these seeding rates was drastically improved. Further, the OT2 dispensing sequence provides control over the exact holding time and the pipette lifting speed away from the scaffold, which may also minimize potential variation.

### 3.5 Single-cell end-point analysis performed on 96-SPOT after automated gel digestion and cell staining

Since the OT2 is designed to transfer liquids, we reasoned that in addition to microtissue manufacturing, the OT2 could assist in other steps that might be useful for high-throughput screening assays. We therefore next set out to optimize a digestion sequence to enable recovery of single cells for downstream end-point analysis, such as using high-throughput flow cytometry. Previous work [22,23,44] using paper scaffolds infiltrated with 5μL of cell-gel showed successful gel digestion and cell retrieval using 1 mL of a solution containing collagenase, protease and DNAase enzymes incubated at 37°C for 45 minutes in a thermomixer, and followed by vigorous pipetting. We applied 200 μL of the same digestion solution with adapted protocols (i.e. no need for a plate shaker during incubation) but focused on optimizing pipetting sequences to be compatible with 96-SPOT digestion. Since the OT2 has an upper limit on liquid dispensing speed to prevent contamination between wells, it is not feasible to achieve the same level of vigorous pipetting as manual pipetting. First, we assessed whether pipetting with the maximum dispensing speed at variable locations (four corners and center) within one well would improve the cell recovery rate compared with the same amount of pipetting at a fixed central location. On day 0, 96-SPOT was seeded with a 5 μL cell-gel mixture and subsequently digested using the OT2. As shown in **Figure 6A**, pipetting at variable locations significantly improved the number of recovered cells. Previously, manual digestion required repeated pipetting 20 times to disrupt the remnant gel and release cells from the paper scaffold. In pilot studies using manual digestion, increasing the enzyme concentration or increasing pipetting results in more significant cell death (data not shown). We tested a range of pipetting times at variable locations within the well and found that repeated pipetting 50 times by the OT2 recovered the highest number of cells among all three pipetting times, which was also the only condition that offered similar recovery levels to the manual pipetting method (**Figure 6B**). More than 30,000 cells were harvested from each well after the washing process, providing a practically useful cell quantity for common downstream analysis.

**Figure 6.**
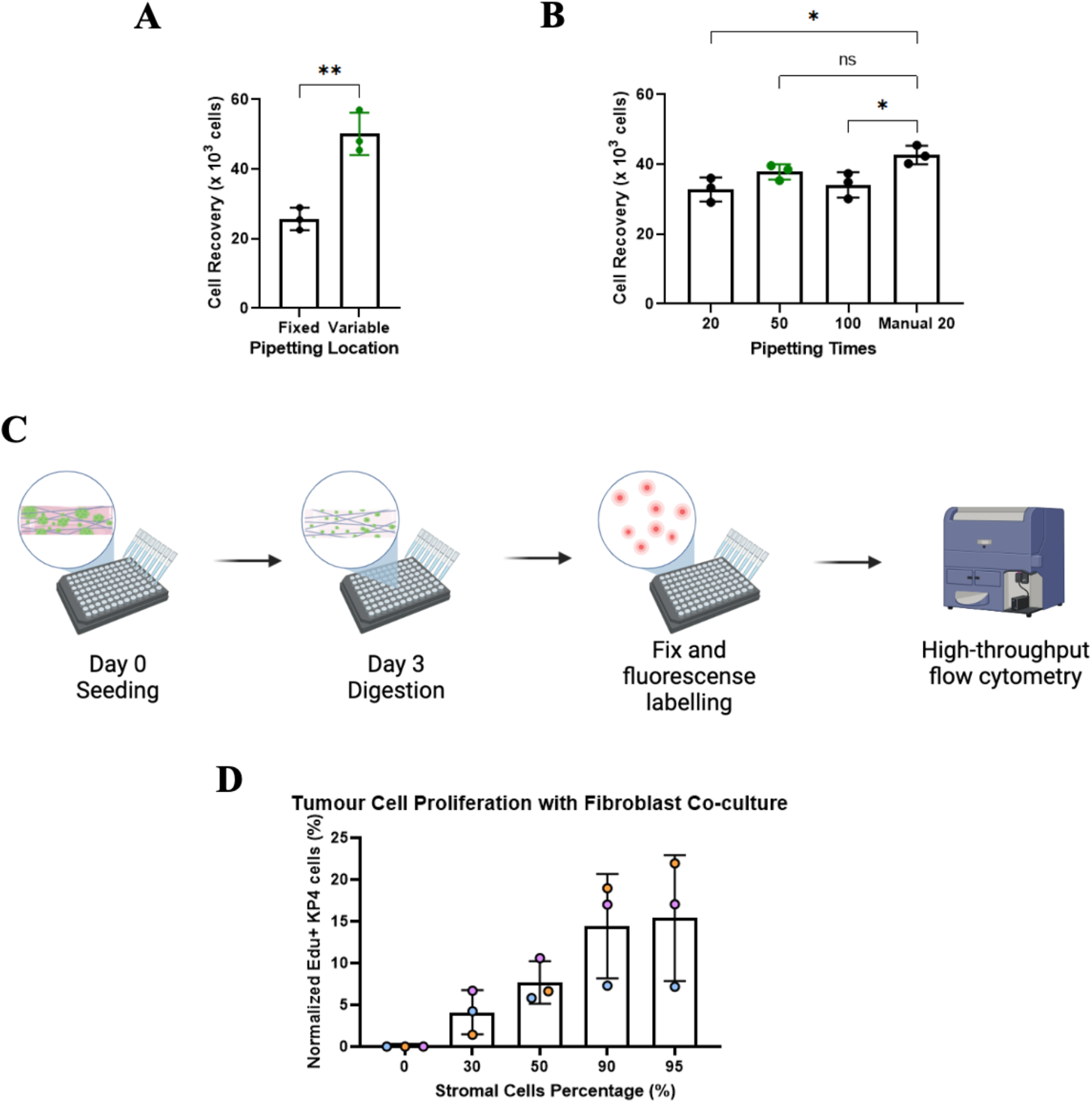
The use of the OT2 to automate cell extraction and facilitate high-throughput end-point single-cell analysis of the 96-SPOT. (A) 30M/mL GFP-expressing KP4 cells were seeded using the OT2 and digested within 2h of seeding. The number of digested whole cells was counted manually by the hemocytometer. Pipetting at variable locations (four corners plus the center) offered a significantly higher cell recovery rate (Student t-test). (B) Fifty times of pipetting offered similar results as manual digestion, while twenty and a hundred times both resulted in significantly lower cell recovery (ANOVA). (C) The timeline and workflow for the proof-of-concept co-culture experiment using KP4 and PSCs. A total cell density of 15 × 10^6^ cells /mL in 6mg/mL bovine collagen was seeded into the 96-SPOT with various tumour and stromal population ratios using the OT2. After three days of co-culture, an EdU assay was performed, and cells were digested out of the paper scaffold by the OT2 for further downstream processing. Analyzed using high-throughput flow cytometry, showed an increasing percentage of proliferative KP4 cells when co-cultured with a higher percentage of PSCs (D). Mean ± SD of 3 biological replicates.

Interestingly, both insufficient and excess pipetting led to a significantly lower cell recovery rate than manual digestion. We speculate this was likely due to the shear stress the cells experienced during excessive pipetting, leading to a smaller number of recovered whole cells. Here, we did not explore the possibility of various digestion formulas, as our main aim was to replace repetitive manual pipetting with an automated sequence specifically for the 96-SPOT with 3mg/mL type I bovine collagen. Similar digestion results and cell recovery rates were achieved as reported in literature using paper scaffold-based models and collagen as matrix [23]. Notably, our paper scaffold is patterned with PMMA instead of wax used in other paper-based platforms [20,21]. It was previously reported that wax-patterned scaffold could introduce particulates during digestion steps [45] which would impose a significant challenge for automating wax-patterned platforms for downstream analysis. In SPOT we did not observed any obvious PMMA debris after digestion.

Having identified a usable automated digestion protocol, we next performed a proof-of-concept experiment to test the entire platform workflow (**Figure 6C**). Previous work has shown that the presence of stromal content promotes the proliferation of cancer cells [46]. Here, we hypothesized that we could also capture this stromal-driven increase in proliferation by co-culturing KP4 cells with pancreatic stellate cells (PSCs) in the 96-SPOT using the OT2 pipeline. Retrieval of cells for single cell analysis is particularly useful in this context because it is impossible to use bulk assays like AlamarBlue™ or Cell-Titer Glo™ to capture changes in a specific population in a co-culture system. As PSCs are more contractive than cancer cells, we used 6 mg/mL of type I bovine collagen for tissue manufacturing to ensure a good distribution of PSC cells after three days of culture (data not shown). Although 6mg/mL type I bovine collagen is slightly more viscous than the 3mg/mL type I bovine collagen that we optimized the OT2 protocols with, no adaptation of the seeding sequences was needed to achieve consistent deposition. After seeding gels containing various stromal and cancer cell compositions, cells were co-cultured for 72h then EdU was added for 3 hours and then digestion of the tissues was performed using the OT2. We also used the OT2 to assist in fixation, washing and the EdU reaction steps to streamline the process and ensure the downstream results were more consistent. Since we used GFP-expressing KP4 and BFP-expressing PSCs in this study, we could easily separate the two populations during flow cytometry and only quantify the number of proliferative cells in the KP4 population. As expected, we observed an increase in GFP KP4 cells proliferation when the ratio of PSCs in the coculture increased (**Figure 6D**). Note that one limitation of this proof-of-concept study was that the digestion and permeabilization protocol was optimized for the extraction and permeabilization of KP4 cells and not the PSC population. Our proof-of-concept study illustrates the possibility of harvesting cells out of paper scaffold using the OT2 for a single cell downstream analysis. We expect our method is compatible with other single-cell-based analyses, such as cytometry-by-time-of-flight and single-cell RNAseq, to assess protein and gene expression on a cellular level. In addition to the whole cell analysis performed here, this protocol could also be useful for bulk assays like qPCR to assess gene expression within each individual well. To perform qPCR, cells must be lysed for RNA extraction and therefore do not require the use of an optimized cell digestion protocol which preserves whole cells. For such an application, this digestion protocol could still be used and then followed with a direct cell lysis protocols to enable recovery of a higher quantity of cell material.

### 3.6 Automated dispensing sequences support the growth of organoid-derived cells

Since prior OT2 optimizations were performed with cell lines, we next wanted to demonstrate the applicability of this workflow to more precious and fragile primary samples, such as patient-derived tumour organoids. To do this, we used a previously tested hydrogel blend of 3mg/mL type I bovine collagen (75%) and Matrigel™ (25%) as the matrix for seeding organoid cells (3× 10^6^ cells/mL of collagen-Matrigel™ blend) into SPOT [17]. Despite the increased temperature sensitivity of the Matrigel™ gel blend, we observed consistent and comparable spreading of cells as measured by the coefficient of variation between manually seeded and OT2 seeded microtissues (**Figure 7A**). Next, we quantified organoid cell growth in SPOT using the AlamarBlue™ assay and the mean gray value of the whole well over 6 days and observed a similar growth trend using both assays (**Figure 7B**). As shown by representative images in **Figure 7C**, GFP-expressing organoid-derived cells exhibited unrestrained growth over time. This further confirmed the capability to track cell growth over time using both high-throughput widefield microscopy and a microplate reader-based assay in the 96/384-SPOT platform (**Figure 7C**). We confirmed organoid cell viability after seeding by performing a dead cell stain using propidium iodide while using the GFP expression as a surrogate for live cells. A similar amount of cell death was observed as with manual seeding, suggesting the decrease in viability after seeding was not caused by the OT2 sequence (**Figure 7D**) but rather likely due to the higher sensitivity of organoid cells compared to cell lines during standard culture manipulations. There was no significant cell death after four days of culture for either seeding method. This was further confirmed by the similar growth trend of organoid cells over 6 days when seeded by the OT2 versus the manual method (**Figure 7E**). Notably, organoid growth from Day 0 to Day 2 was less prominent than at later time points. We speculate that this was likely because the organoid cells were recovering from the seeding process which involves digestion from full organoid structures into single cells and incubation on ice until further seeding. Thus, we expect that before using these microtissues for any drug screening assays, 2 to 3 recovery days should be given. Significantly, the OT2 seeding sequence was compatible with these precious primary samples without requiring significant modifications. While we only tested one pancreatic ductal adenocarcinoma (PDAC) organoid line, we speculate that our workflow is likely appropriate for PDAC organoid originating from different patients or organoids originating from other cancer types or from healthy tissues with a similar matrix composition to facilitate comparable wicking seeding.

**Figure 7.**
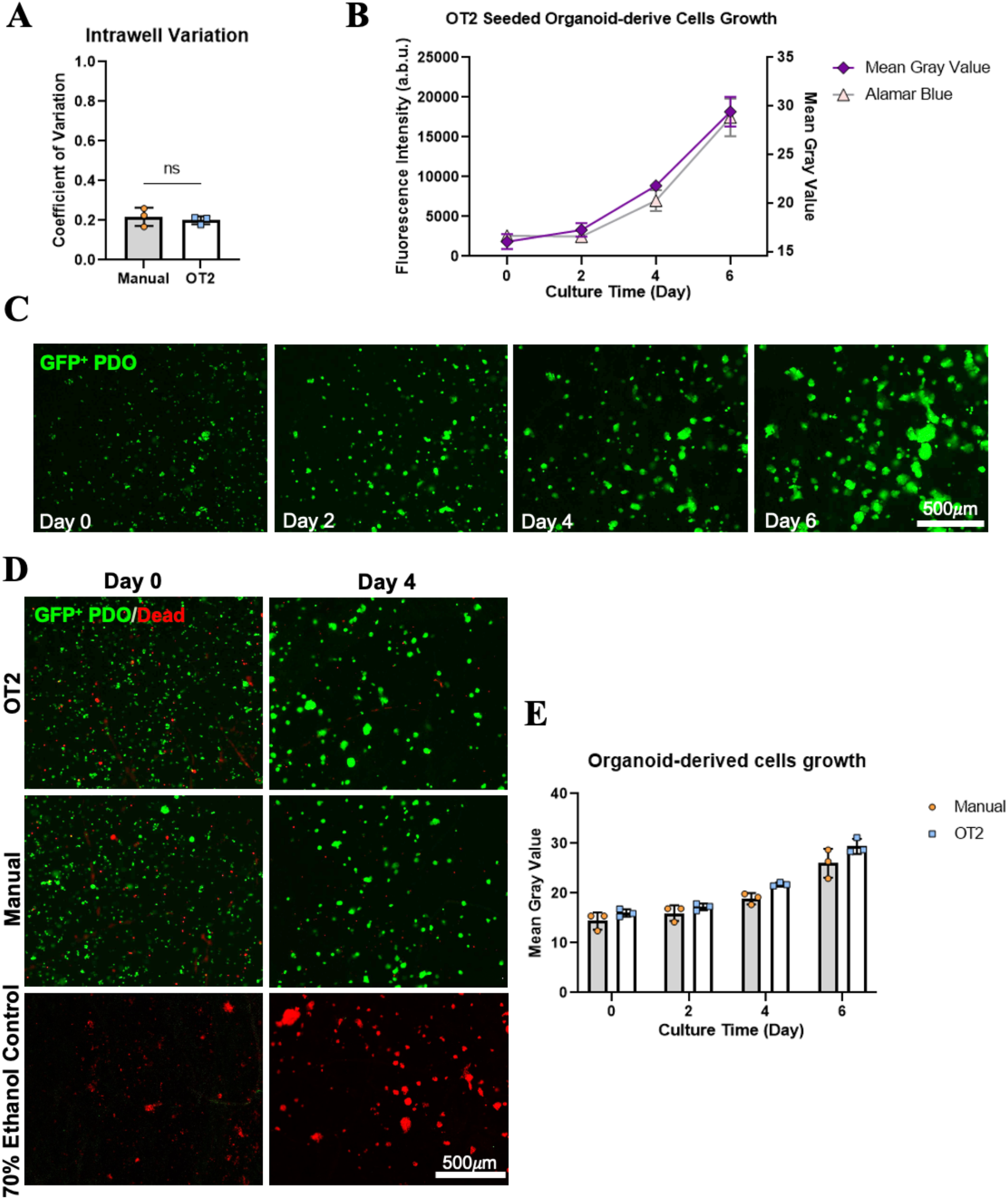
Translation of automated OT2 pipeline to incorporate fragile patient-derived organoid cells (A) Assessment of intrawell variation between OT2 and manual seeded wells on day 0. GFP-expressing PPTO.46 cells were seeded at 3 × 10 ^6^ cells/mL in a hydrogel blend (3mg/mL bovine collagen (75%) and Matrigel™ (25%). No significant differences were observed (revealed by Student t-test for 3 independent experiments). Plot shows mean ±SD of 3 independent experiments. (B) Representative widefield fluorescent images of SPOT microtissues containing GFP-expressing PPTO.46 cells (green) grown over six days. (C) Growth curve for GFP-expressing PPTO.46 cells after being seeded using the OT2 in SPOT comparing measurements of the imaging-based readout (right-hand axis, mean gray value) and plate-reader-based assay (left-hand axis, AlamarBlue™). (D) Representative images of GFP expressing PPTO.46 seeded using either the OT2 or manual on Day 0 and Day 4. Dead cells (red) were stained with propidium iodide, while GFP expression was used as the surrogate for live cells (green). Cells were treated with ice-cold 70% ethanol for 10 minutes to induce complete cell death. (E) Growth curve for manually seeded and OT2 seeded microtissues. Mean gray values of the GFP signals were obtained over 6 days of growth. Plots show the mean ±SD of 3 independent experiments. ANOVA revealed no significant differences between cells seeded manually or using the OT2.

In this report we describe the automation of both 96 and 384-SPOT. We envision that 384-SPOT could serve as a platform for medium to high-throughput screening relying on time-course imaging analysis or plate-reader-based assays (such as AlamarBlue™ and Cell-Titer Glo™) as metrics to distinguish hits. On the other hand, 96-SPOTcontains sufficient cellular material to offer the possibility for single-cell-based end-point analysis to decipher specific gene expression or analyze based on cell types for example in a co-culture model. Notably, the established seeding pipeline is optimized based on a P20 8-channel pipette; thus, the number of distinct tissue compositions that can be generated within each plate is limited. In the future it could be possible to incorporate a single channel pipette for cell-gel deposition, however, the run time could be eight times longer; thus, it might not be ideal for cell survival, particularly in fragile primary samples. Thus, the most optimal application for the current pipeline is likely using a few distinct tumour compositions per plate. The use of conditioned media from other cell types is also possible to add another layer of complexity to the microenvironment. This would allow the optimized OT2 downstream analysis protocols, rapid and consistent single-cell-based analysis to be used to decipher the effect of specific tumour microenvironment effects on a subpopulation of cells. Further, given we have shown our approach is compatible with primary patient samples, we envision applications to investigate patient-level heterogeneity in therapy response. This could potentially open a new avenue for screening patient-specific drug targets in a high-throughput but biologically relevant system.

We note that this report describes in detail our steps to optimize the automation of SPOT manufacturing and analysis for pancreatic tumour cells in a collagen matrix blend specifically. While alterations in cell type or matrix will likely require slight adjustments to the parameters we selected here, our work provides a starting point and highlights key parameters we identified that impacted manufacturing quality. Further, the described cell digestion protocols will likely need further optimization depending on the specific cell types and matrix being used. Interestingly the OT-2 workflow developed here was compatible with all three different viscous hydrogel compositions we used without the need for protocol adjustments. There is considerable effort within the community to generate biomatrix alternatives to replace commonly used Matrigel™ since it cannot be easily standardized [47,48]. Many of these alternative bio-matrices are viscous hydrogels or hydrogel precursors therefore automated tissue manufacturing methods compatible with these more viscous materials will likely be important as the use of these alternative gels increases [49]. We anticipate our workflow is likely compatible with many of these synthetic gels, particularly those with a thermal gelation step, however some parameter optimization may be necessary following similar steps to those described here.

## 4. Conclusion

In this study, we present an automated pipeline using a commercially available liquid handler to streamline all the steps from the fabrication of 3D tissue arrays to their analysis. By combining the use of the SPOT platform and the OT2 liquid handler, we eliminated labour-intensive steps while preserving the consistency and robustness of fabricated tissues across wells and plates. Beyond executing plate-reader-based assay to assess bulk response, this automated workflow allowed us to probe the effect of tumour microenvironment cues on specific cell types through high-content automated microscopy and high-throughput flow cytometry at a single-cell level. Furthermore, we demonstrated that this workflow supports the incorporation of patient-derived primary samples with minimal adaptation. This highlights the potential application of this model for investigating patient-level heterogeneity to assist in personalized medicine discovery and clinical management. Further, the workflow takes advantage of an open-source and inexpensive OT2 liquid handler, to allow for easy implementation into existing drug discovery pipelines and research facilities with the possibility of combining with big-data analysis in the future. We envision that our work will offers new avenues for discovering novel personalized medicines.

## Acknowledgements

The authors acknowledge technical assistance from Saifedine T Rjaibi, Ileana Co, Erik Jacques, Nila C Wu, and Xiaoya Lu. Figures 1, 3B and 6C were created with BioRender.com.

## Data and materials availability

The datasets generated during and analyzed in this study are available from the corresponding author upon reasonable request.

