## Supplementary Information for "An automation workflow for high-throughput manufacturing and analysis of scaffold-supported 3D tissue arrays"

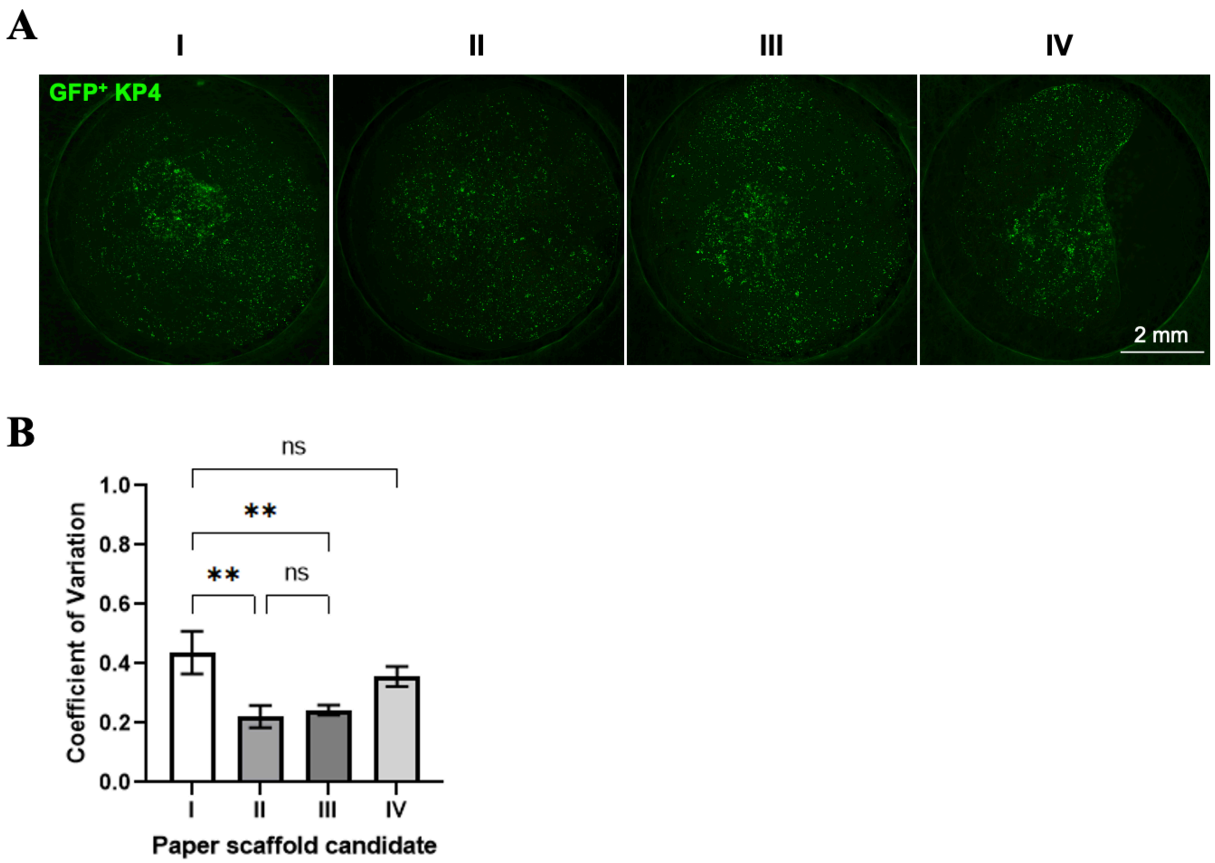


**SI Figure 1.** Comparison of tissue homogeneity seeded on all four scaffolds. (A) Representative widefield images of a complete 96-SPOT well, showing scaffold IV leads to less homogeneous tissues. (B) The variation within each well represented by the coefficient of variation metric obtained from widefield fluorescent images of GFP-expressing KP4 cells infiltrated in each scaffold candidate at a cell density of 5 x $10^{6}$ cells/mL type I bovine collagen hydrogel. Scaffold II and III offered significantly lower coefficient of variation than scaffold I, which indicates a better cell-gel mixture spreading within the well. Statistical significance was assessed using ANOVA. Mean + SD of 3 independent experiments. Scalebar is 2 mm.

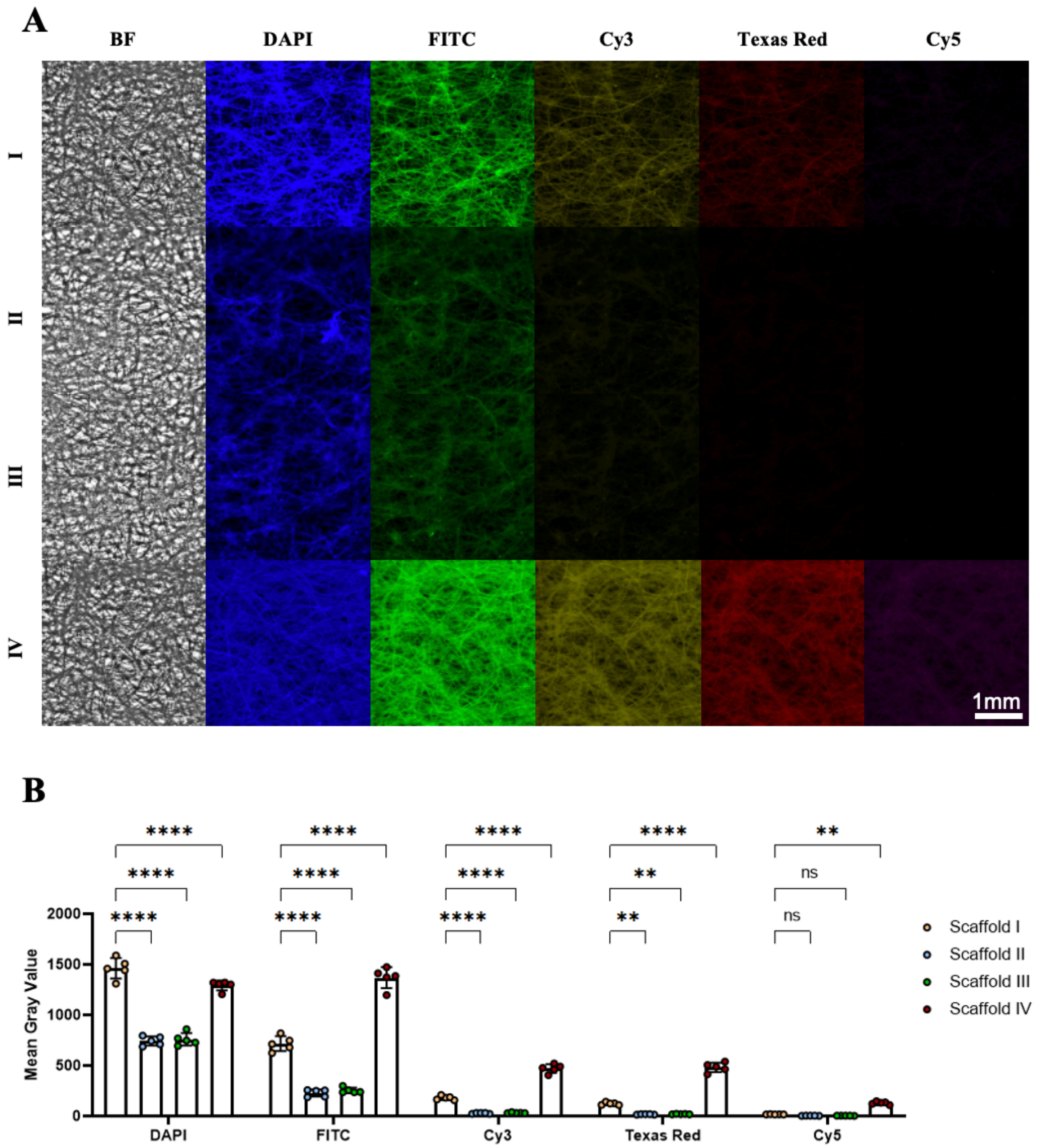


**SI Figure 2.** Paper scaffolds autofluorescence characterization. (A) Representative widefield images showing brightfield (BF, gray), and autofluorescence of 4 paper scaffold candidates (I to IV, respectively, from top to bottom) in the DAPI (blue), FITC(green), Cy3(yellow), Texas Red (red) and Cy5(magenta) channels, respectively from left to right. Each fluorescent image was taken with an exposure time of 300 ms in order to compare the autofluorescence directly. (B) Quantification of autofluorescence based on mean gray value for each fluorescence channel and scaffold. Scaffold II and III emit significantly lower autofluorescence than the original scaffold in DAPI, FITC, Cy3 and Texas Red channels, while scaffold IV has much stronger autofluorescence in all 5 channels. Scalebar is 1mm.
